# Nonlinear compliance of NompC gating spring and its implication in mechanotransduction

**DOI:** 10.1101/2024.06.20.599842

**Authors:** Yukun Wang, Peng Jin, Avinash Kumar, Lily Jan, Yifan Cheng, Yuh-Nung Jan, Yongli Zhang

**Affiliations:** Department of Cell Biology, Yale University School of Medicine, New Haven, CT, USA; Department of Physiology, University of California, San Francisco, CA, USA; Department of Biochemistry and Biophysics, University of California, San Francisco, CA, USA; Howard Hughes Medical Institute, UCSF, San Francisco, CA, USA; Department of Molecular Biophysics and Biochemistry, Yale University, New Haven, CT, USA

## Abstract

Cytoskeleton-tethered mechanosensitive channels (MSCs) utilize compliant proteins or protein domains called gating springs to convert mechanical stimuli into electric signals, enabling sound and touch sensation and proprioception. The mechanical properties of these gating springs, however, remain elusive. Here, we explored the mechanical properties of the homotetrameric NompC complex containing long ankyrin-repeat domains (ARDs). We developed a toehold-mediated strand displacement approach to tether single membrane proteins, allowing us to exert force on them and precisely measure their absolute extension using optical tweezers. Our findings revealed that each ARD has a low stiffness of ∼0.7 pN/nm and begins to unfold stepwise at ∼7 pN, leading to nonlinear compliance. Our calculations indicate that this nonlinear compliance may help regulate NompC’s sensitivity, dynamic range, and kinetics to detect mechanical stimuli. Overall, our research highlights the importance of a compliant and unfolding-refolding gating spring in facilitating a graded response of MSC ion transduction across a wide spectrum of mechanical stimuli.

---

Mechanosensitive ion channels (MSCs) detect mechanical forces impinging on cell membranes arising from touch, sound, blood and osmotic pressure, cell motion, etc.^1,2^ While most MSCs sense the force within membranes or membrane tension, other channels are tethered to extracellular or intracellular structures via flexible filaments or gating springs and detect the force out of membranes^3,4^ (Fig. 1a). One example of the latter is the MSC in hair cells of the inner ear, which is essential for hearing^5,6^. However, these tethered MSCs are poorly understood, partially due to the challenges in reconstituting the MSC systems *in vitro* and applying precise force to MSCs. NompC/TRPN is a representative of these tethered MSCs. It belongs to the transient receptor potential (TRP) family channels and is found in various sensory organs in a wide range of organisms from worms to vertebrates, where it mediates sound and touch sensation and proprioception^7–9^. NompC is a homotetramer. Each subunit is mainly composed of a pore-forming transmembrane domain and a cytosolic ankyrin repeats domain (ARD, Fig. 1b) that binds the microtubule^8,10^. Previous studies showed that the ARD plays a pivotal role in conveying force from the cytoskeleton to the channel^8^. But the details of force transmission and gating process remain elusive The mechanical properties of gating springs play crucial roles in mechanosensation by tethered MSCs^11^. The gating springs could bear significant force even in a rest cell state and experience large force changes in response to mechanical stimuli, necessitating their mechanical stabilities^12^. These stimuli are represented by displacements of membranes, cytoskeleton, hair bundles, and other sensors ranging from subnanometers to hundreds of nanometers^13^. To transduce such a large range of displacement to typically nanometer conformational changes in the transduction channel, a flexible gating spring is required to confer a gradual current response. Furthermore, the stiffness and stability of the gating spring need to be regulated to balance the sensitivity and dynamic range to detect the mechanical stimuli^12^. The sensitivity to small displacements requires a high gating-spring stiffness, whereas large displacements need a low stiffness or protein unfolding^14^. Consequently, the stiffness of a gating spring is expected to be nonlinear with force.

**Fig. 1.**
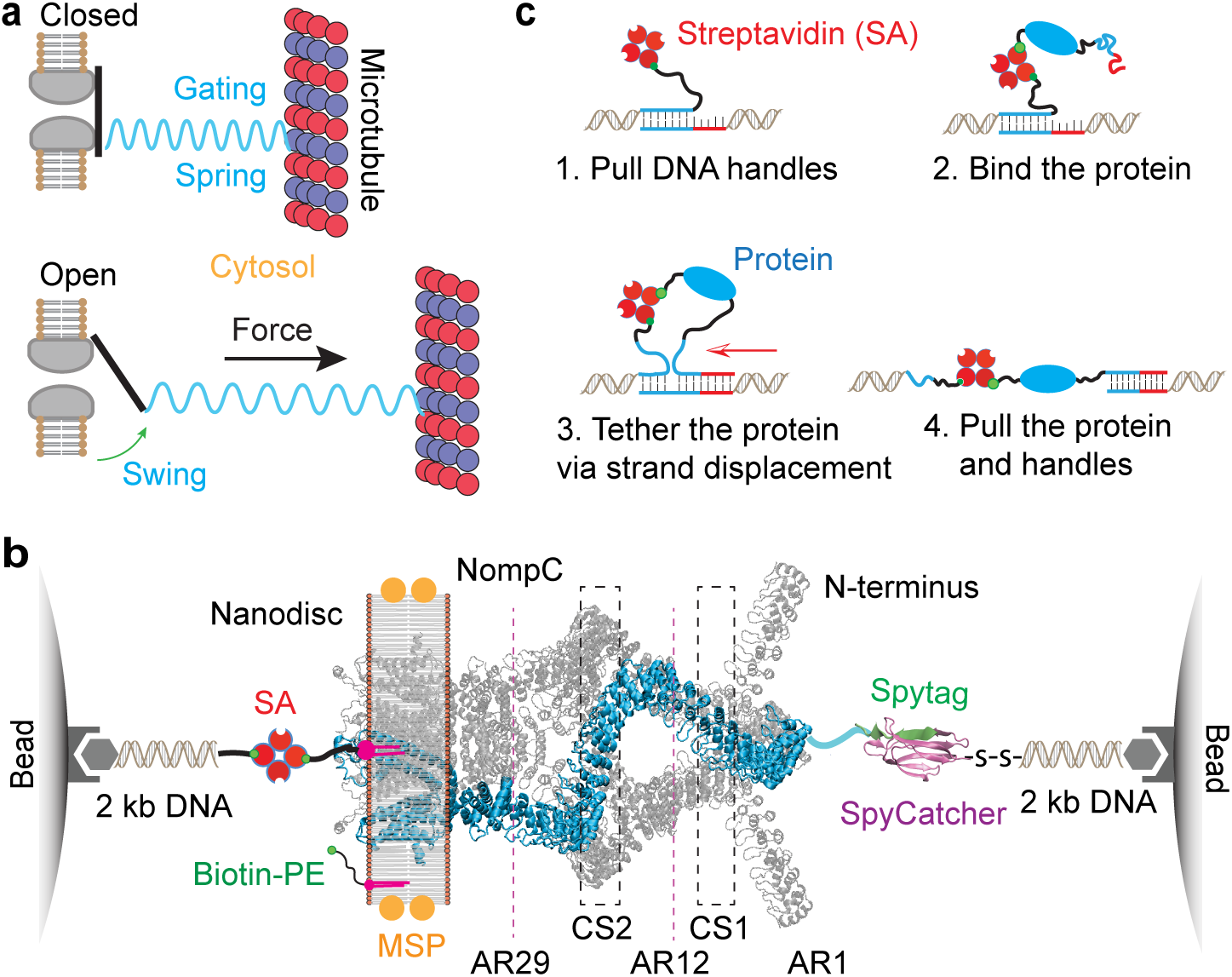
Toehold-mediated strand displacement strategy to measure the elasticity and unfolding of a single NompC complex using high-resolution optical tweezers. **a.** Schematic diagram showing the channel opening transition of NompC induced by the force exerted by the microtubule and transmitted by the gating spring. **b.** Experimental setup to pull a single NompC homotetramer embedded in a lipid nanodisc (PDB ID 5VKQ). The NompC complex was tethered between two 2.1 µm beads via two DNA handles through biotin-streptavidin (SA) and Spytag-SpyCatcher interactions. Positions of ankyrin repeats (AR1-29) and the two ARD contract sites (CS1 and CS2) are shown. MSP, membrane scaffolding protein; SA, streptavidin. **c.** Steps to link a single protein to two pre-stretched DNA handles via toehold-mediated strand displacement and pull the protein using dual-trap optical tweezers (see also Movie 1).

The stiffnesses and mechanical stabilities of ankyrin repeats are crucial for their functions but are not well understood. The 33-amino acid ankyrin repeat is one of the most abundant motifs whose tandem repeats occur in over 400 proteins in humans^15^. Many ARDs link the cytoskeleton to diverse membrane targets. Howard and Bechstedt proposed that the NompC ARDs act as a flexible gating spring and estimated a stiffness of ∼ 1 pN/nm per ARD based on the helical structure of the ARD and the shear modulus of rigid proteins^16^. Similarly, the 24 ankyrin repeat domain in the ankyrin repeat B protein (AnkB) may also serve as part of the gating spring of C. elegans TMC1^6^, whose mammalian homologs TMC1/2 have been generally considered as the long-sought MSC responsible for hearing^6,17^. However, the AnkB ARD appears to be much stiffer than the NompC ARD, as atomic force microscopy (AFM) revealed that the AnkB ARD has a stiffness greater than 1.9 pN/nm^18,19^. Molecular dynamics simulations yielded an even higher stiffness of 4-5 pN/nm for the AnkB ARD^20^. Likewise, molecular dynamics simulations revealed a stiffness of 4 pN/nm for a single NompC ARD^21^ or 13 pN/nm^22^ for the four ARDs together as the NompC gating spring, both of which are substantially higher than the earlier estimation. In addition, AFM experiments showed that AnkB ARD was mechanically stable, with a high unfolding force above 20 pN^18,19^, while the mechanical stability of NompC ARD has not been measured. As the AFM measurement is limited by the low resolution for force detection (∼10 pN) and the nonspecific pulling site on the protein^18^, a more accurate method is required to characterize the mechanical properties of ARDs to provide insights into how forces gate MSCs.

Tethered MSCs generally undergo discrete conformational changes to open their channels, which are associated with energy increases^13^. These energy increases are compensated by the mechanical work done by the gating force, which necessitates the corresponding extension changes, or gating swing, in the direction of the force application (Fig. 1a). Therefore, the gating swing is a projection of the conformational changes in the direction of the gating force. Both gating swing and force are characteristic parameters of tethered MSCs and critical for their sensitivity and dynamic range to detect mechanical stimuli. For the mammalian hearing channel in hair cells, the gating swing is estimated to be 2-6 nm^13,23^, while the gating force ranges from 5 pN to 35 pN, depending upon the cell type, or inner or outer hair cells, and their locations in the cochlear^12^. However, the gating swing and gating force of NompC have not been reported.

We developed a toehold-mediated strand displacement strategy to measure both monotonic and discrete extension changes of a single NompC complex in response to force through high-resolution optical tweezers^24,25^. Using this strategy, we characterized the mechanical properties of single NompC complexes. We found that individual ARD is compliant and unfolds at a low force, which may facilitate the transmission of mechanical stimuli to ion transduction of the channel domain. We also observed a conformational change of NompC potentially correlated to its gating transition. Our results corroborate the proposed role of the ARDs as a gating spring for NompC and shed light on its gating mechanism.

## Results

### A differential approach for measuring the absolute extension of a single protein complex using optical tweezers

Despite their wide applications to soluble proteins, optical tweezers have rarely been used to characterize the dynamics of membrane proteins at a single-molecule level, partly due to a lack of model membranes compatible with optical trapping^26–28^. We generated a construct expressing NompC with a SpyTag attached to its N-terminus^29^, purified and reconstituted the protein into a lipid nanodisc containing biotin-labeled phosphatidylethanolamine (PE)^10,30^. A single NompC tetrameric complex in the nanodisc was connected to two optically trapped polystyrene beads (∼2 microns in diameter) via two DNA handles, one conjugated to the N-terminus of one subunit of NompC and the other to the biotin-PE lipid in the nanodisc (Fig. 1b). These ∼2 kbp DNA handles are essential for attaching a single macromolecule to micron-sized beads and accurately measuring force and extension measurements by optical tweezers^31^. We stretched one single NompC subunit or monomer by moving one optical trap away from the other and measured the extension and tension of the whole NompC-nanodisc-DNA complex tethered between the two beads.

To differentially measure the absolute extension of a single macromolecule in the presence of a pulling force, we developed a method named toehold-mediated strand displacement strategy. Despite their subnanometer resolution to detect discrete relative extension changes from a single tether between two beads, high-resolution optical tweezers have far less resolution, typically greater than 10 nm, to measure the absolute extensions of different tethers^25,32^. The measured tether extension is sensitive to variations of bead sizes and DNA attachment sites on bead surfaces, which cause reduced resolution to measure the absolute extensions of both the protein-DNA tether and the protein alone, as the latter is derived from the former. The reduced resolution makes it difficult, if not impossible, to detect the elasticity of a single protein. Our new strategy overcame the issue by measuring the absolution extension of a single protein using the same DNA tether attached to the same pair of beads (Fig. 1c & Movie 1). First, we attached a DNA-only tether between two beads, in which two DNA handles were joined in the middle through the hybridization of two 5’-overhang DNA sequences (Fig. 1c, state 1). Then, we pulled the DNA tether to measure its force-extension curve (FEC, Fig. 2a, grey curve), leading to an FEC that was well fit by the worm-like chain model of DNA molecules^33,34^ (blue dashed curve). Next, we relaxed the DNA tether and held it at a low force when a NompC solution flowed into the microfluidic channel. A single NompC complex bound to one of the overhang DNA sequences via biotin-PE in the nanodisc (Fig. 1c, state 2). The NompC complex was attached with a DNA oligonucleotide at the SpyTag through the SpyTag–SpyCatcher interaction. With part of its sequence identical to the hybridized DNA overhang, the oligonucleotide displaced the DNA overhang via toehold-mediated strand displacement^35^ (Fig. 1c, state 3), as manifested by a sudden extension decrease (Fig. 2a, red arrow). This strategy allows us to accurately measure the absolute extension of a single protein by subtracting the extension of DNA handles from the extension of the whole DNA-protein tether.

**Fig. 2.**
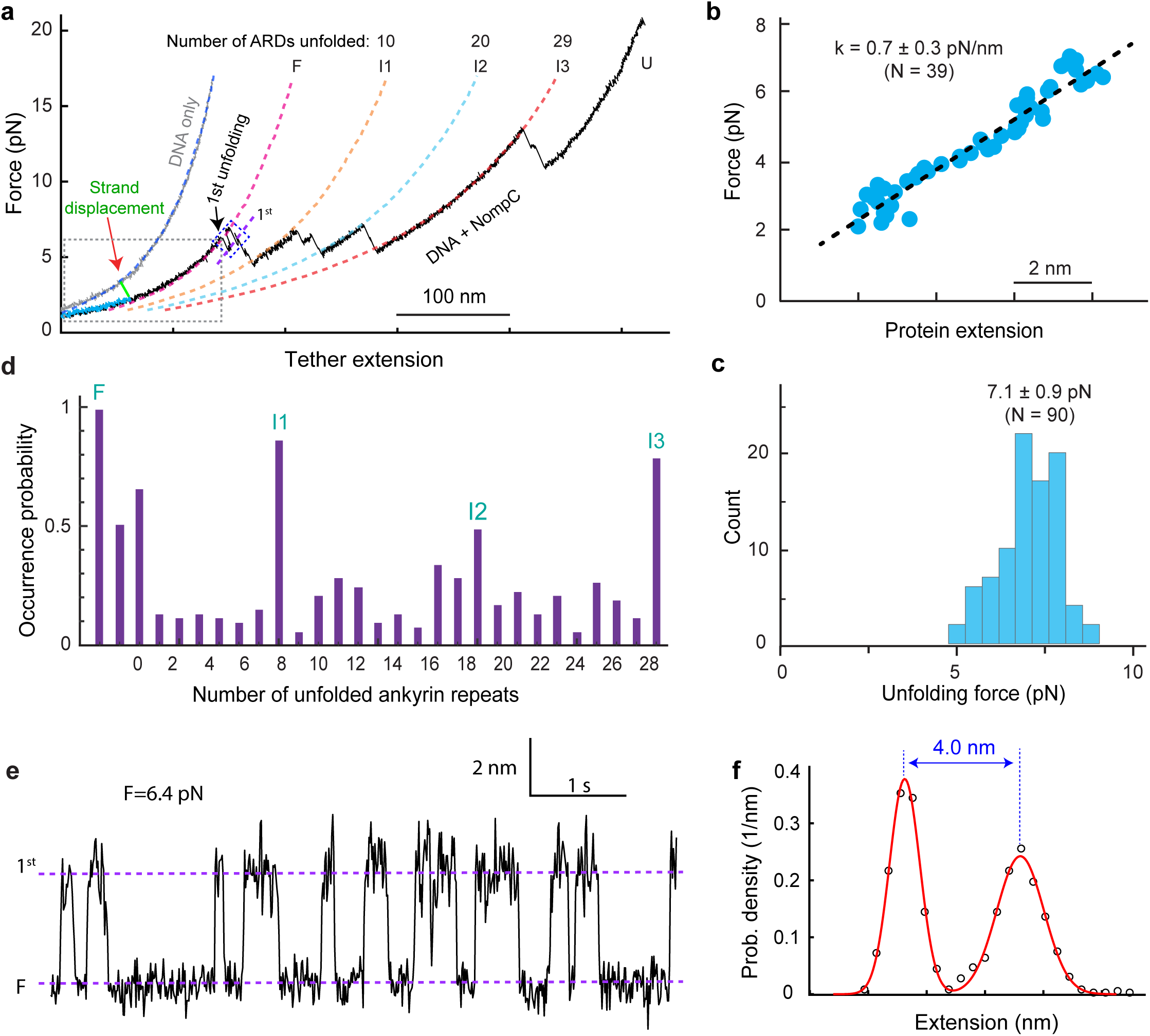
Mechanical properties of a single NompC complex. **a.** Representative force-extension curves (FECs) for pulling DNA handles only (gray) and the NompC-DNA tether (black). Different states are indicated: F, the folded native state; U, the cytosolic domain fully unfolded state; and I1-3, the intermediate unfolding states. The nanodisc-embedded NompC was inserted into the DNA handles by the strand displacement (green FEC). The dashed gray rectangle marks the FEC regions used to calculate the force constant of the gating spring (see **c**). The dashed lines are bet-fits of different FEC regions with a worm-like chain model, showing the numbers of ankyrin repeats unfolded in different major intermediates (I1, I2, I3, and U). **b.** Force as a function of the extension of NompC only (symbol) and its linear fit (dashed line) to derive the stiffness of the protein. **c.** Histograms of the force for the first abrupt conformational transition of NompC (with one event marked in **a**). **d.** Occurrence probabilities of different unfolding states derived from the FECs. The three long-lived states I1, I2, and I3 appear as three peaks. **e.** Extension-time trajectory at a constant mean force (F) corresponding to the first reversible transition (region marked by the dashed blue rectangle in a). The dashed lines indicate the average extensions of the native NompC state (F) and the first intermediate state (1^st^). **f.** Probability density distribution of the extension shown in e (circle) and its best fit with a sum of two Gaussian functions (red curve).

### Elasticity and unfolding of the NompC complex

When pulled with forces up to ∼7 pN, the NompC-nanodisc-DNA tether first continuously increased in extension, due to elastic stretching of the NompC complex and the DNA handles (Fig. 2a). After subtractions, we plotted the force as a function of the extension of NompC only (Fig. 2b). The NompC extension increases linearly with its extension, revealing a force constant of 0.7 (± 0.3, SEM) pN/nm. The force corresponding to the first unfolding event varied from 5 pN to 8 pN, with an average of 7.1 (± 0.9, S.D.) pN (Fig. 2c). Therefore, a NompC complex can extend ∼10 nm on average before unfolding. In contrast, globular proteins are generally brittle and extend no more than a few nanometers before unfolding by force^31^. The comparison demonstrates that single NompC complexes are exceptionally elastic. Above the force threshold, successive discrete extension increases or flickering occurred (Fig. 2a), indicating irreversible or reversible conformational transitions, respectively (Fig. 2d).

When different NompC complexes were subjected to forces larger than 10 pN, their FECs overlapped, indicating that the entire cytosolic domain or the ARD of the NompC monomer was unfolded (Fig. 2a, state U; Extended Data Fig. 1, state I3). The assignments of different unfolding states were confirmed by their different contour lengths or numbers of residues in the unfolded polypeptides derived from fitting of the FEC regions to a worm-like chain model (see Method). Despite their stochastic nature, the FEC regions corresponding to various unfolding intermediates showed consistent patterns among different NompC complexes being pulled (Fig. 2d), which reveals more than 30 unfolding intermediates (Extended Data Fig. 1). Three of these states stand out due to their long lifetimes and high occurrence frequency (states I1, I2, and I3 in Figs. 2a, 2d and Extended Data Fig. 1). Their extensions relative to the folded and unfolded states indicate that the NompC ARD likely unfolded from the N-terminus to different degrees – roughly one-third for state I1 (∼10 repeats unfolded), two-thirds for state I2 (∼20 repeats unfolded), and fully unfolded for state I3 (all 29 repeats unfolded). However, more work is required to validate these proposed ARD folding states corresponding to the intermediates. Thus, the NompC ARD readily unfolds in a stepwise manner with distinct intermediates.

### Refolding properties of the NompC complex

To investigate whether the unfolded ARD could refold, we relaxed the tension applied to the unfolded NompC subunit. When the forces reached below 6 pN, the protein started to refold (Fig. 3a, discrete extension decreases indicated by red arrows). When force was below 1.5 pN, the NompC complex returned to an extension close to its native state, leading to hysteresis between the pulling and relaxation FECs (Fig. 3a, compare cyan and black curves). Pulling the refolded NompC again revealed a FEC that significantly overlaps the FEC from the first pulling round (Fig. 3a, compare green and black FECs for NompC #1), which further supports that this NompC complex refolded into its native state. More experiments revealed that ∼30% of unfolded NompC complexes could successfully refold (Fig. 3b). This fraction of successful NompC refolding under our experimental conditions is surprisingly high, given NompC’s large size and intricate structure. The remaining refolded NompC complexes showed FECs that significantly differ from that of native NompC complexes (Fig. 3a, NompC #2, compare green and black traces), indicating that these NompC complexes were misfolded. To further pinpoint NompC refolding, we relaxed the NompC complex from less unfolded I2 and I1 intermediates to test their refolding probabilities (Extended Data Fig. 2). We observed that the NompC complexes refolded with progressively higher probability from less unfolded intermediates (Fig. 3b). Particularly, the intermediate I1 refolded with a high probability of 94% under our experimental conditions. Thus, the ARDs of the NompC monomers readily unfold at a relatively low force and could refold from unfolding intermediates with a high probability when relaxed to a low force.

**Fig. 3.**
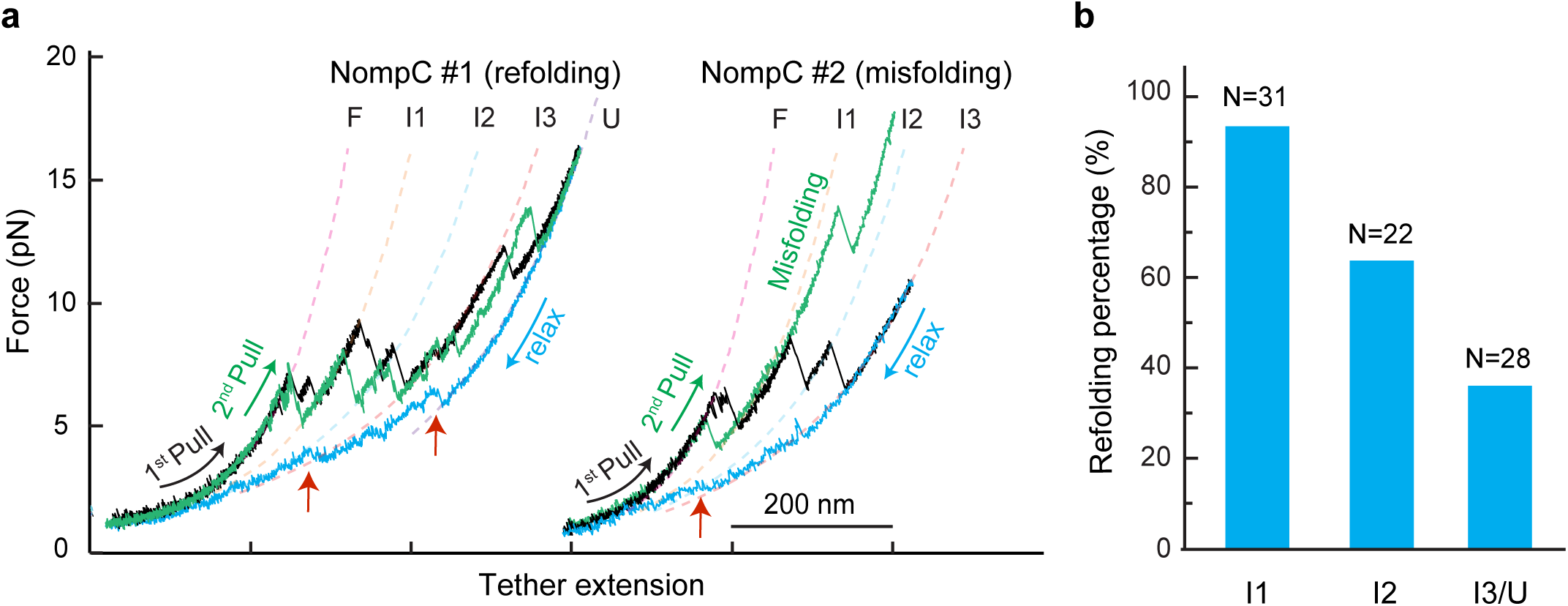
The unfolded ARDs could refold and misfold. **a.** FECs for pulling and relaxing a single NompC complex for multiple consecutive rounds showing NompC refolding (NompC #1) and misfolding (NompC #2). The black and green FECs are for the first and second rounds of pulling, respectively, while cyan FECs are for relaxation. **b.** Probabilities of successful NompC ARD refolding from three major intermediates.

### The first reversible transition as a potential gating transition

Interestingly, the first intermediate underwent reversible transition with the folded native NompC state before further unfolding in ∼60% pulling FECs (Fig. 2a, between state 1^st^ and state F & Extended Data Fig. 1, regions marked by dashed blue rectangles). In the remaining pulling FECs, the first intermediate probably unfolded without sufficient time to visit the native state. The reversible transition was also observed when NompC was held at a constant mean force (Fig. 2e). The transition is two-state as indicated by the bimodal distribution of the probability density function of the extension, which can be well fit by a sum of two Gaussian functions (Fig. 2f). The average equilibrium force for the transition, i.e., the force with equal probability in either state, was 7.0 ± 0.8 (S.D., N=10) pN, while the corresponding extension change was 4.6 ± 0.7 nm. This led to an energy change of 8.0 ± 1.5 k_B_T or 4.7 ± 0.9 kcal/mol. We hypothesize this reversible transition may represent the gating transition of NompC, with more evidence described in the forthcoming sections.

### Mechanical properties of AnkB ARD

To investigate how the mechanical properties of a single NompC subunit depend on the other three subunits in the complex, we pulled a single AnkB ARD and compared its mechanical properties with those of NompC ARD described above^15^ (Fig. 4a). A single isolated ARD domain would be a better choice for the comparison than the AnkB ARD. Unfortunately, we failed to purify the isolated ARD due to its high propensity to aggregation. Nevertheless, the purified AnkB ARD is structurally intact when examined under an electron microscope (Fig. 4b & Extended Data Fig. 3). We revisited the mechanical properties of this ARD using high-resolution optical tweezers (Fig. 4c & Extended Data Fig. 4). To our surprise, we found that the AnkB ARD began to unfold at a lower average force of 5.5 (± 0.7, S.D.) pN (Fig. 4d) and had a much lower force constant of 0.12 ± 0.03 (SEM) pN/nm (Fig. 4e), compared with the NompC ARD. Interestingly, the AnkB ARD unfolded via at least 23 intermediates with comparable occurrence frequency (Fig. 4f), suggesting highly parallel ARD unfolding pathways. Finally, the AnkB ARD always refolded to its native state with little hysteresis upon relaxation (Fig. 4g). Previous AFM measurements of the same AnkB ARD yielded significantly higher stiffness (>1.9 pN/nm) and unfolding force (>20 pN) than our measurements^18,19^. However, our findings are consistent with previous observations that the IκBα ARD (composed of six ankyrin repeats) is marginally stable at room temperature even in the absence of force^36^. The differences between our results and the AFM measurements probably resulted from the higher force resolution offered by high-resolution optical tweezers (∼0.02 pN compared with ∼10 pN for AFM)^24,37^, the differential measurement via the strand displacement strategy, and specific pulling sites on the ARD. Taken together, our measurements on the mechanical properties of ARDs in NompC and AnkB demonstrate that ARDs are generally compliant, unfold at a relatively low force, and can readily refold. Their significant differences in stiffness and unfolding imply strong interactions between the four ARDs within a single NompC tetramer (Fig. 5a). As one subunit is pulled, the lateral interactions transmit force to other subunits, likely through the two major contact sites among ARDs (CS1 and CS2 in Fig. 1b)^10^. This results in an increased force constant for the entire ARD superhelix compared to isolated individual ARDs. Interestingly, we did not observe a similar reversible transition for AnkB ARD as we did for NompC (Fig. 4c). This finding is consistent with our hypothesis that the first reversible NompC transition may be a gating transition, which is expected to be independent of ARD unfolding.

**Fig. 4.**
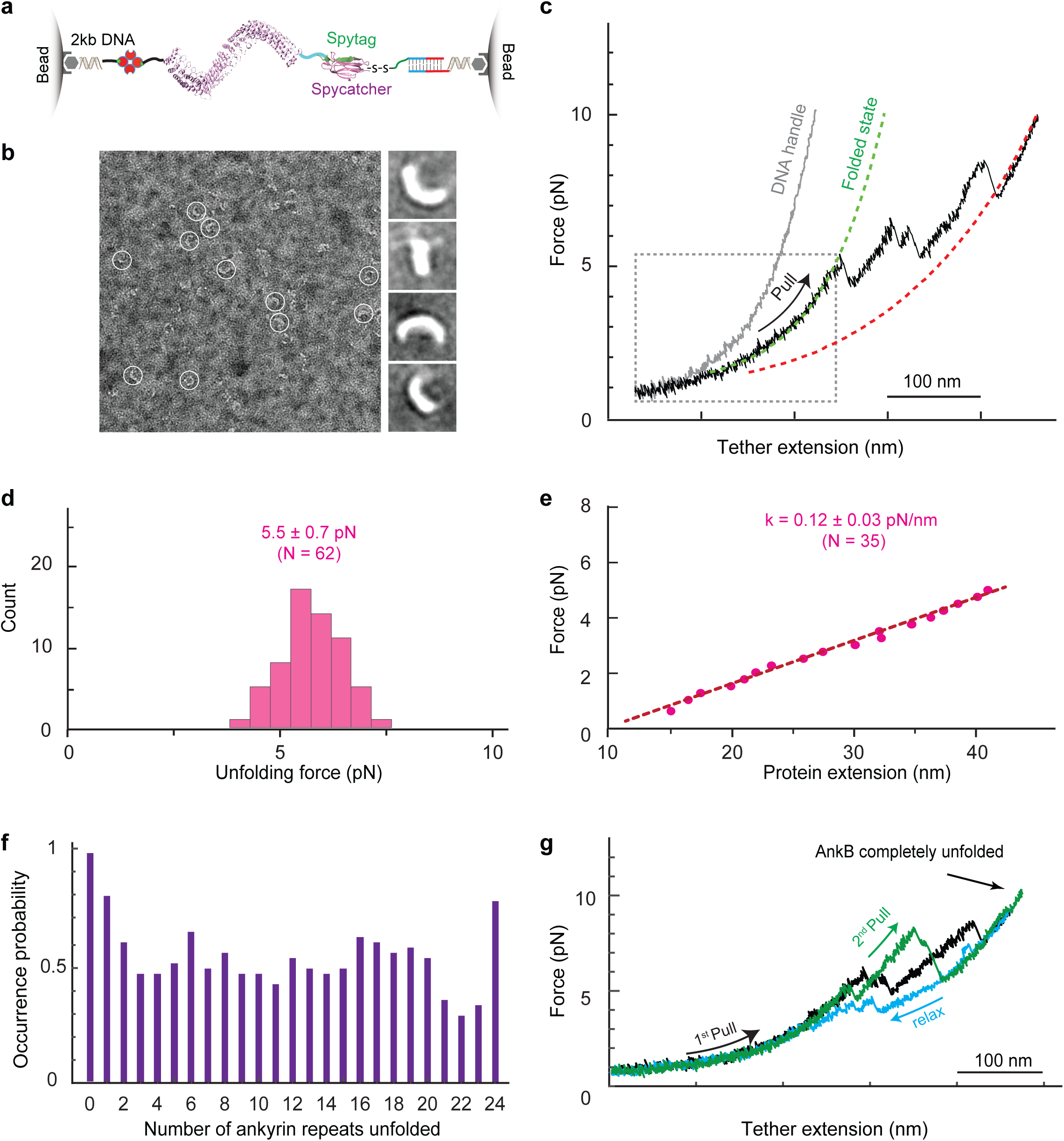
Compliant and refoldable AnkB ARD. **a**. Schematic diagram illustrating the attachment of a single AnkB ARD to polystyrene beads using two DNA handles. **b**. EM images of purified AnkB ARD revealed its expected elongated helical shape. **c.** FECs obtained by pulling the DNA handle alone (gray) and the protein-DNA tether (black) using the toehold-mediated strand displacement strategy. **d.** Histograms of the force for the first unfolding transition of AnkB ARD. **e.** Force as a function of the extension of AnkB ARD (symbol) and its linear fit (dashed line) to derive the stiffness of the protein. **f**. Occurrence probabilities of different unfolding states derived from the FECs. **g**. FECs obtained by pulling (black and green) and relaxing (cyan) a single AnkB ARD for multiple rounds show efficient ARD refolding.

**Fig. 5.**
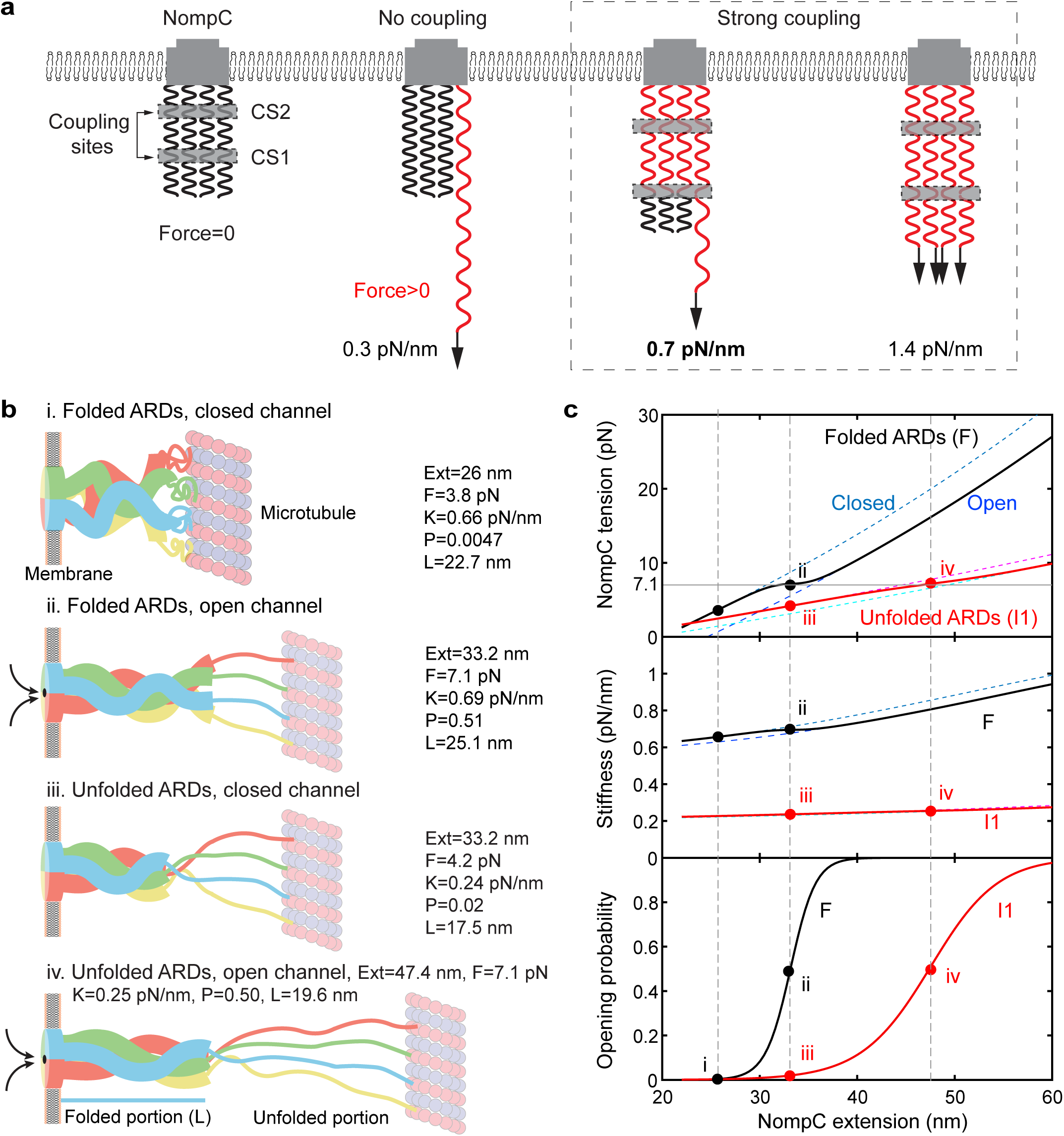
Theoretical calculations highlight the roles of ARD mechanics in modulating force-triggered channel opening. **a.** The two contact sites (CS1 and CS2) joined the four ARDs into a coupled spring with an effective force constant depending on the number of ARDs being pulled. Each ARD is modeled as a uniform elastic rod with its compliance proportional to its length. The stretched and relaxed ARD regions are shown in red and black, respectively. The stiffnesses shown were derived from the stiffness measured (in bold) by pulling a single NompC subunit (see Data analysis and modeling in Method). **b.** Schematics of characteristic ARD unfolding and channel states at different NompC extensions. The NompC extension (ext), tension (F), stiffness (K), channel opening probability (P), and the extension of the structured portion (L) associated with these states are indicated by their values, as well as their positions in **c** indicated by dots. **c.** NompC tension (top panel), stiffness (middle), and channel opening probability (bottom) as a function of the NompC extension. The fully folded and partially unfolded ARD states are indicated by black and red curves, respectively, while the open and closed channel states are plotted by nearby dashed curves.

#### Role of the nonlinear compliance in channel gating

To explore the biological significance of the NompC mechanics we discovered, we derived the tension and stiffness of the gating spring and the channel opening probability of the entire NompC tetramer mimicking the cellular context (see Data analysis and modeling in Method for details). Besides the assumption for the gating transition, our model additionally assumed that the four NompC subunits extend cooperatively in response to the pulling force (Fig. 5a). Cryo-electron tomography analysis of NompC in fly haltere revealed that NompC complexes are overstretched to a broad range of extensions from 20 nm to 80 nm, with an average extension of ∼43 nm, or more than double the 20 nm length of the NompC structure^10,38,39^. To accommodate such a large extension, NompC ARDs are likely partially unfolded *in vivo*^38^. Therefore, we modeled the folded portion of NompC ARD tetramer as an elastic rod with the derived stiffness of 1.4 pN/nm and their unfolded polypeptide chains as worm-like chains^33^ (Fig. 5b). For simplicity, we only considered the folded ARD state (F) and the partially unfolded state (I1) in our calculations, with all four subunits bound to the microtubule. In the fully folded NompC, the 123 residues at the N-terminal end are disordered in each subunit^8,10^ (Fig. 5b, states i & ii), while the partially unfolded state contains a total of 442 disordered residues (states iii & iv). The calculated tension of a single NompC tetramer in a folded ARD state is higher in a closed channel state than in an open state, and both increase with the NompC extension (Fig. 5c, top panel, cyan and blue dashed curves). The average tension (black curve) gradually changes from the tension of the closed channel state at a low force to the tension of the open channel state at a high force, with a middle turning point at 7.1 pN. Given the same NompC extension, the partial ARD unfolding reduces the tension (compare red and black curves). The stiffnesses of the NompC tetramers are generally lower than that of the folded gating spring (1.4 pN/nm, Fig. 5c, middle panel), as the stiffness of the unfolded polypeptide is significantly smaller than that of the folded portion. To simulate the impact of the gating spring on channel opening, we chose the gating swing of 4.6 nm and the gating transition energy of 8 k_B_T for the NompC channel derived by us. The calculated opening probability of the channel is sigmoidal, with a gating force of 7.1 pN for a half-channel-opening probability (Fig. 5c, bottom panel). The probability strongly depends on the folding states of the gating spring and shifts to a higher NompC extension as the gating spring unfolds (compare the red curve to the black one). The slope of the sigmoidal distribution at a half-opening probability represents the sensitivity of NompC to detect mechanical stimuli. Thus, a folded gating spring confers a higher sensitivity but a lower dynamic range than a partially unfolded gating spring for NompC to detect mechanical stimuli. This observation is justified by the fact that the former has significantly greater stiffness than the latter (Fig. 5c, middle panel).

Our model can explain several prominent features of NompC channel kinetics observed *in vivo*. First, the sudden ARD unfolding in response to a large and prolonged displacement reduces the force exerted on the channel, causing it to close (Figs. 5b & 5c, from state ii to state iii). Thus, the ARD unfolding could underlie NompC inactivation or insensitivity to small mechanical stimuli immediately after strong ones. Second, the unfolded ARDs allow the NompC channel to adapt to larger mechanical stimuli with reduced sensitivity, causing a right shift of the channel opening probability as observed by electrophysiological measurements^40,41^ (Fig. 3b, bottom panel). Third, slow ARD refolding in the absence of large mechanical stimuli restores the stiffness of the gating spring and sensitivity to the small stimuli (from state iii to state i to state ii). Finally, highly dynamic gating springs may be essential for tethered MSCs in general, which act as an autonomous gain switch to allow their channels to detect a large range of mechanical stimuli. This is exemplified by tip links, which are gating springs for the mammalian hearing channel and also undergo force-dependent unfolding/refolding or association/dissociation^14,42–44^.

## Discussion

In this study, we measured the mechanical properties of NompC using a new differential approach to detect the absolute extension of a single molecule with improved accuracy based on optical tweezers. We found that ARDs in NompC and AnkB are much more compliant than previous estimations or measurements. We also observed ARD unfolding and refolding in a physiological force range and explored its role in channel gating. It remains a matter of debate as to whether pulling or pushing force triggers the opening of the NompC channel. Molecular dynamics simulations and experiments on cell surfaces imply a pushing or compression force gates the NompC channel^21,22^. Particularly, the pushing force generates a torque to twist open the channel. However, it is unclear how the torque is applied to the transduction channel without a mechanism to constrain its free rotation in the membrane. Like many elastic objects, the stiffnesses of NompC gating spring measured by us are expected to be independent of pushing or pulling force at least in a low force range. However, ARD unfolding induced by pushing force, if it occurs, should be different from that caused by pulling force.

The mechanical properties of NompC are comparable to those reported for the hearing channels in hair cells. The force constant of the NompC gating spring ranges from 0.2 pN/nm to 0.9 pN/nm, while the force constant of the gating spring for the hearing channel ranges from 0.4 pN/nm to 1.8 pN/nm in inner hair cells^12^. In addition, the gating swing (4.6 nm) and force (∼7 pN) of the NompC channel derived by us match those reported for the hearing channels in hair cells (∼4 nm and 3-35 pN)^12–14,27^. However, it must be noted that the first reversible transition observed by us may be affected by the small size of the nanodisc and its association with the gating transition needs to be validated by its correlation to the ion transduction through the channel. Finally, the gating springs in both NompC and the hearing machinery may undergo reversible unfolding and unbinding to modulate channel kinetics^14^. Since NompC is also involved in sound detection in some species, the comparisons suggest that the two tethered MSCs may detect mechanical stimuli through shared molecular mechanisms.

Further experiments will focus on incorporating single-channel recording into the current single-molecule assay to directly dissect the gating mechanism^45^.

## Method

### Expression and purification of NompC

*Drosophila* full-length NompC was expressed and purified using the BacMam system as previously described (Jin et al., 2017). In brief, an N-terminal Spytag, the gene encoding NompC, and a C-terminal 3C cleavage site with eGFP were assembled through Gibson assembly into a modified pFastBac vector with a mammalian CMV promoter. Bacmids were extracted from DH10Bac *E. coli* cells after the transformation of the NompC construct. Subsequently, the bacmid was transfected into *Spodoptera frugiperda* (Sf9) cells to generate baculoviruses. The protein was expressed in HEK293 GnTi^-/-^ suspension cells and grown in ESF SFM Serum-Free Medium (Expression Systems LLC), supplemented with 1% FBS (Axenia BioLogix) at 37 °C. The culture was supplemented with 10 mM sodium butyrate (Sigma) to boost protein expression 24 hours after baculovirus infection and then further incubated at 30 °C for 48 hours before harvest.

The NompC complex was affinity purified at 4 °C using GFP nanobody. The cell pellet from 0.5 L culture (∼3 g) was first resuspended and lysed in 50 mL hypotonic buffer containing 10 mM Bicine pH 8.5, 1 mM EDTA, 3 mM dithiothreitol (DTT), 1× complete protease inhibitor cocktail (Roche) and 1 mM phenylmethylsulfonyl (PMSF). The membrane fraction was collected by centrifugation at 35,000 x g for 30 min and then dispersed again by homogenization in extraction buffer (20 mM Bicine pH 8.5, 500 mM NaCl, 1 mM EDTA, 1x complete protease inhibitor cocktail, 1 mM PMSF). Protein was extracted in 50 mL extraction buffer supplemented with 1% n-Dodecyl-β-D-Maltopyranoside (DDM) and 0.2% Cholesteryl hemisuccinate (CHS) with a gentle stir. After a two-hour extraction, the lysate was cleared by centrifugation at 35,000 x g for one hour. The supernatant was mixed with 1 mL CNBr-activated sepharose resin (GE Healthcare) coupled with an anti-GFP nanobody (Kirchhofer et al., 2010) and incubated on a rotator overnight. The resin was then loaded onto a polypropylene column and washed with 10x column volume of wash buffer (WB) containing 20 mM Bicine pH 8.5, 500 mM KCl, 0.025% DDM, 0.005% CHS, 1 mM EDTA, 0.05mg/ml 3:1:1 1-palmitoyl-2-oleoyl-sn-glycero-3-phosphocholine (POPC):1-palmitoyl-2-oleoyl-sn-glycero-3-phosphoethanolamine (POPE): 1-palmitoyl-2-oleoyl-sn-glycero-3-phospho-(1’-rac-glycerol) (POPG). The washed resin was incubated quiescently with 1 mL WB containing 0.2 mg 3C precision protease overnight to release the protein.

### Reconstituting NompC into nanodiscs

The concentration of the eluted protein was determined based on its absorbance at 280 nm. Soybean polar lipid extract (Avanti) was prepared in water as described previously (Jin et al., 2017). For structural study by EM, the protein sample was mixed with MSP2N2 and soybean lipids at a molar ratio of NompC monomers:MSP2N2:soybean lipids =1:2:100 (molar ratio). Additional functional lipid was added before reconstitution for optical tweezers experiments.

Biotin-PE (1,2-distearoyl-sn-glycero-3-phosphoethanolamine-N-[biotinyl(polyethylene glycol)-2000]) was incorporated as soybean lipids:Biotin-PE =7:1 (molar ratio) for pulling the NompC complex. The protein-lipid mixture was put on constant rotation till reconstitution was complete. After a two-hour incubation, Bio-Beads SM2 (Bio-Rad) were supplemented to the mixture three times at over 3-hour intervals to initiate reconstitution by gradually removing detergents from the system. Each time 20 mL of briefly dried Bio-Beads were added per 1 mL protein sample used for reconstitution. One day after the last batch of Bio-Beads, the beads were removed by passing the sample through a mini-column, and the cleared sample was applied to a Superose-6 size exclusion column equilibrated with CB. The peak fractions corresponding to NompC were pooled and immediately shipped at 0 °C through Fed-Ex for the optical tweezers experiments.

### Expression and purification of SpyCatcher

The plasmid of SpyCatcher was generated by mutagenesis of SpyCatcher002 plasmid (purchased from Addgene with Plasmid #102827), which added three amino acids (KCK) to the C terminus of the encoded protein sequence. The expression and purification of SpyCatcher were conducted as previously described^46^. Briefly, the protein was transformed into Escherichia coli BL21 (DE3) cells, grown in LB broth (Miller) at 37 ℃ until OD_600_ reached 0.6-0.8, and expressed by adding 0.4 mM IPTG at 30 ℃ for 4-5 h. The cells were harvested by centrifuging the culture at 5000 × g, resuspended in the lysis buffer (50 mM Tris-HCl, 300 mM NaCl, 1 mM DTT, pH 7.8) with 1 × EDTA-free protease inhibitor cocktail (cOmplete™), and clarified by centrifugation at 35000 × g for 1 h at 4 ℃. The resulting supernatant was mixed with pre-washed Ni-NTA resin (Cytiva) overnight at 4 ℃ and washed with the lysis buffer containing 10, 20, and 40 mM imidazole. The SpyCatcher protein was eluted from the resin in the lysis buffer containing 300 mM imidazole, dialyzed into a buffer containing 50 mM Tris-HCl, 300 mM NaCl, 1 mM TECP, and 5% glycerol, pH 7.8, and finally aliquoted and stored at −80 °C before use.

### Expression, purification, and biotinylation of AnkB ARD

The original plasmid of Ankyrin-B ARD was provided by Mingjie Zhang’s group^15^, which contains the gene coding for residues 28-872 of human AnkB cloned into the pDEST14 vector. The protein construct was modified by adding a Spytag003 tag (RGVPHIVMVDAYKRYK)^46^ to the N-terminus of the ARD and an Avi-tag (GLNDIFEAQKIEWHE) to the C-terminus. The plasmid was transformed into Escherichia coli BL21 (DE3) cells for protein expression. The purified AnkB ARD was biotinylated at the Avi-tag using the BirA biotin-protein ligase (Avidity) in the imidazole-free lysis buffer at 4 ℃ overnight. The biotinylated protein was dialyzed in 4 L buffer containing 50 mM Tris, 200 mM NaCl, and 1 mM EDTA, pH 7.5, to remove free biotin and DTT.

### Characterization of the structural integrity of AnkB ARD

Protein quality was examined by size exclusion chromatography through a superdex-200 column (GE health care) as well as negative stain EM. Negative stain EM was performed as described previously (Booth et al., 2011). Images were acquired on a Tecnai T12 microscope (FEI Company) operated at 120 kV and a nominal magnification of 52,000× using an UltraScan 400 camera (Gatan), corresponding to a pixel size of 2.21 Å on the specimen. 2D averages of AnkB ARD particles were analyzed by EMAN2 (Tang et al., 2007).

### Oligo-SpyCatcher

The oligo-SpyCatcher linked the protein to be pulled to one of DNA handles (Figs. 1c and 4a). The chemically synthesized oligonucleotide was cross-linked to the C-terminus of SpyCatcher through a disulfide bridge and has the sequence: 5’-GAGGGCGTACAGTTGTATGTACGTTGGCGAGTTT – SH.

This oligo could hybridize to the DNA handle with a complementary overhang sequence. To crosslink the oligo to SpyCather, the oligo was first treated with 2 mM tris(2-carboxyethyl)phosphine (TCEP) and then buffer-exchanged into a low pH Buffer A (100 mM NaH_2_PO_4_, 400 mM NaCl, pH 5.8) using Micro spin-bio chromatography columns. Next, 2 mM 2,2’-dithiodipyridine disulfide (DTDP) solution was added to the oligonucleotide solution and incubated at room temperature for 2 h. Excess DTDP was removed by exchanging Buffer A for Buffer C (100 mM NaH_2_PO_4_, 400 mM NaCl, pH 8.5). The SpyCatcher solution was similarly exchanged to Buffer C, mixed with DTDP-treated oligonucleotides with 1:50 protein-to-oligo molar ratio, and kept overnight at room temperature to allow disulfide bond formation between oligos and SpyCatcher. The oligo-labeled SpyCatcher was bound to Ni-NTA resin via the his-tag on SpyCatcher and rotated at 4 ℃ for 2 h, extensively washed with Buffer C, and eluted with a buffer containing 300 mM KCl, 20 mM bicine, 300 mM imidazole, pH 8.3.

### Preparation of DNA handles

Two DNA handles, designated as left and right handles (Fig. 1c), were used in our experiments. They have the same length of 2,260 bp and dual digoxigenin labels at one end but different overhang oligonucleotide sequences at the other end. The two DNA handles were made by polymerase chain reaction (PCR) with different templates and primer sets (ordered from Eurofins Genomics) listed below:

Left DNA handle:

Forward primer: [DIG]-ATCATCCAA-[DIG]-GGCTGAGCCTGCAGG

Reverse primer: <u>[Biotin]-TTTTTTTTAAGTATGTACGTTGGCGAG</u>-**S18**-AAATCGACGCTCA AGTCAGA

Right DNA handle:

Forward primer: [DIG]-TCGCCACCA-[DIG]-TCATTTCCAGCTTTTGTG

Reverse primer: <u>CTCGCCAACGTACATACAACTGTACGCCCTC</u>-**S18**-ACTATCGCCACTT TTATTGGCG

Here S18 indicates the 18-atom hexa-ethylene glycol spacer used to prevent polymerase extension to the overhang regions (underlined sequences) during PCR.

### Dual-trap high-resolution optical tweezers

Our optical tweezers system is based on a custom-made microscope, constructed on a vibration-isolation table (TMC, MA), within a temperature-controlled room. The optical tweezer is operated in a dual-trap mode, with one trap fixed and the other movable. Detailed specifications can be found elsewhere^25^. In brief, a 1064 nm laser beam (CW, Nd: YVO4, Spectra-Physics, 5 W) is expanded by a 5X telescope, collimated, and split (50:50) using a polarized beam splitter (PBS). The two split beams with orthogonal polarizations are reflected by a fixed mirror and a mirror mounted on a high-resolution piezoelectric actuator (Mad City Labs, WI) and combined with another PBS. The combined beams are further expanded by a 2X telescope and focused by a water-immersion, high-numerical-aperture objective (Olympus 60X, NA = 1.2) to trap two 2 μm diameter anti-digoxigenin coated polystyrene beads (DIGP-20-2, Spherotech Inc.) inside a microfluidic chamber^47^. The microfluidic chamber contains three channels where the top and bottom channels flow beads to the middle channel, where they are trapped. The piezo-actuated mirror can turn along two axes and thus move one of the optical traps in the sample plane. To detect displacements of the trapped beads, the outgoing trapping beams are collected by a second identical microscope objective, split by polarization, and projected onto two position-sensitive detectors (Pacific Silicon Sensor). An LED light source and a CCD camera are used to illuminate and view the microfluidic chamber, respectively. A LabVIEW interface is used to control and collect data from optical tweezers. To calibrate each optical trap, the Brownian motion of the trapped bead is measured. The power spectral density of the corresponding displacement trajectory is calculated and fit with a Lorentzian distribution to derive the trap stiffness and voltage-to-displacement conversion factor^47^.

### Single-molecule experiments

All pulling experiments were performed using dual-trap high-resolution optical tweezers as previously described with the adaption for toehold-mediated strand displacement strategy. 2 μL 0.4 μg/μL left DNA handle was mixed with 2 μL 1 mg/ml streptavidin and incubated at room temperature for 10 minutes. Then 2 μL of 100-fold dilution of the DNA-streptavidin solution and an aliquot of 10 ng right DNA handle were each mixed with 10 μL polystyrene beads 2.1 μm in diameter coated with anti-digoxigenin antibody (DIG beads, Spherotech) and incubated at room temperature for 10 min. Next, the DIG beads were diluted in 1 mL working buffer (20 mM Bicine pH 8.5, 300 mM KCl). Subsequently, the two bead solutions were separately injected into the top and bottom channels of the microfluidic chamber^47^. The central channel contained the working buffer with an oxygen scavenging system comprising 10 mg/mL glucose (Sigma-Aldrich), 0.02 unit/mL glucose oxidase (Sigma-Aldrich), and 0.06 unit/mL catalase (Sigma-Aldrich). A single left DNA handle-bound DIG bead was trapped and brought close to a single right DNA handle-bound DIG bead held in another optical trap to form a single ∼4 kbp DNA tether between the two beads.

To pull the NompC complex, the nanodisc-embedded NompC complex was mixed with oligo-SpyCatcher with a 2:1 molar ratio and incubated at 4 °C overnight to allow conjugation of the oligo-labeled SpyCatcher to the NompC complex. Then the NompC complex was diluted to ∼4 nM in the working buffer and flowed into the central channel, where the single DNA tether was being held at a constant force. Binding of the NompC complex to the DNA tether triggered the toehold-mediated strand displacement, leading to the attachment of the NompC complex to the two DNA handles. The flow rate was kept constant at 10 μL/min with a flow control system. AnkB ARD was pulled similarly.

The stiffness of NompC or AnkB ARD was measured by a toehold-mediated strand displacement strategy. The DNA-only tether was first pulled to measure its force-extension curve (FEC). Then, the DNA tether was relaxed and held at around 5 pN force, waiting for the protein solution being injected into the central channel. Protein insertion into the DNA tether via strand displacement was identified by a sudden decrease in the tether extension. Finally, the protein-DNA tether was pulled or relaxed to measure the corresponding FECs. All pulling and relaxation were conducted by moving one trap at a speed of 10 nm/sec while the position of the other trap was kept fixed.

### Data analysis and modeling

We modeled the unfolded polypeptide and the DNA handle with a worm-like chain model for a semi-flexible chain^33^. The stretching force *F* and the entropic energy *E* of the polymer chain are related to its extension *x*, contour length *L*, and persistence length *P* by the following formulae

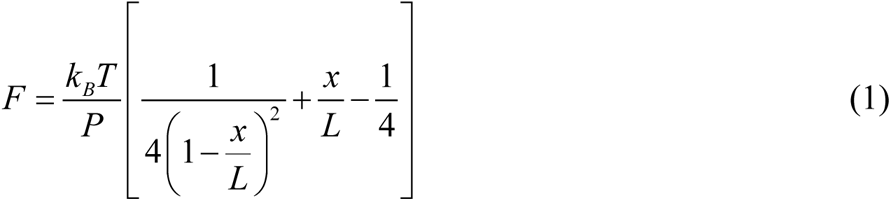

and

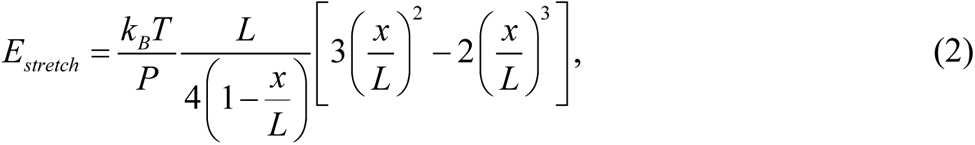

respectively. We adopted a persistence length of 40 nm for DNA and 0.6 nm for the unfolded polypeptide^48–50^. The contour length of a polypeptide was calculated from its number of amino acids with 0.365 nm per amino acid. The extension of a protein *x_P_* consists of the contributions from its folded or structured portion *x ^f^* and the unfolded portion *x_uf_* . The folded portion of the protein *x _f_* is modeled as an elastic rod with stiffness *k* and an intrinsic length *x*_0_ at zero force. The extension of the unfolded portion *x_uf_* was modeled by Eq. (1). Therefore, the extension of a single protein as a function of its tension *F* can be expressed as

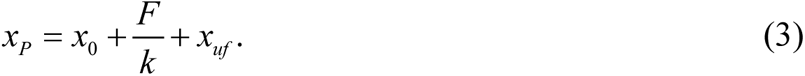

The extension of a protein-DNA tether *x_PD_* is the sum of the extensions of the protein and the DNA handle *x_DNA_*, or

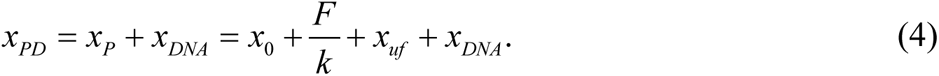

The force-dependent extension (FEC) of the protein was calculated as the difference between the extensions of the DNA-only tethers and the protein-DNA tethers. This extension was subtracted by the extension of the unfolded portion of the protein to yield the extension of the folded protein portion. Its extension was fit with a straight line to derive the protein stiffness *k* and the intrinsic length *x*_0_ (Figs. 2b & 4e). To fit the FECs shown in Fig. 2a, the FECs of the DNA-only and protein-DNA tethers were simultaneously fit by Eqs. (1) and (4) for *x_DNA_* and *x_PD_*, respectively, with the persistence lengths for polypeptide and DNA and the intrinsic length and stiffness of the folded protein portion as fitting parameters.

To calculate the tension *F* of NompC given its extension *x_P_*, we solved the Eq. (3) for *F* . The intrinsic extension of the structured NompC portion *x*_0_ varies with ARD folding and channel states. In a closed channel state, *x*_0_ = 20 nm for the folded ARDs and *x*_0_ = 14.5 nm for the partially unfolded ARDs, while in an open channel state, a gating swing of 4.6 nm was added to the intrinsic extensions. The compliance (the inverse of the stiffness) of the entire NompC gating spring is the sum of the compliances of the folded and unfolding portions of the protein, with the latter calculated from Eq. (1). The opening probability of the NompC channel was determined by the Boltzmann distribution of the closed and open states. The total energy of a stretched NompC *E* was the sum of the elastic energy of the folded portion of the gating spring, the entropic energy of the unfolded portion *E_uf_*, and the channel energy *V*, or

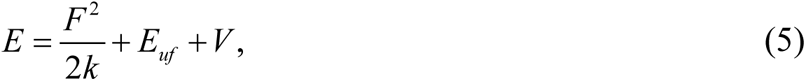

where the entropic energy *E_uf_* was calculated using Eq. (2) and *F* was the tension. The three terms in Eq. (5) depend on the channel state and the ARD folding state, with *V* = 0 for the closed channel state and *V* = 8 k_B_T (k_B_T=4.1 pN×nm) for the open state. The channel opening probability was computed based on the Boltzmann distribution, i.e.,

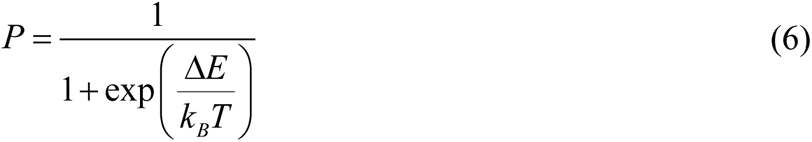

where Δ*E* is the energy difference of NompC in the open state and closed state.

To estimate the stiffnesses of NompC with multiple subunits being pulled (Fig. 5a), we assumed that each NompC ARD has uniform elasticity along its length and four subunits within a single NompC complex are linked at the N-terminal contact site (CS1), or the 9^th^ ankyrin repeat from the N-terminus. Suppose each isolated ARD with 29 ankyrin repeats has a stiffness of *k_a_*, then the two ARD portions separated by CS1 have stiffnesses of 29*k_a_ /*9 and 29*k_a_ /*20 for the N-terminal portion and the C-terminal portion, respectively. The C-terminal portions of four ARDs act as four springs in parallel in response to pulling force and have an effective stiffness of *k_C_* = 4 × 29*k_a_*/20 = 29*k_a_*/5. The N-terminal portions of ARDs also act in parallel but have an effective stiffness *k_N_* with a spring number equal to the number of subunits being pulled (*N*), or *k_N_* = *N* × 29*k_a_ /*9. Since the N-terminal and C-terminal portions of ARDs are connected in series, the overall stiffness of the NompC complex as a function of the subunits being pulled can be calculated as

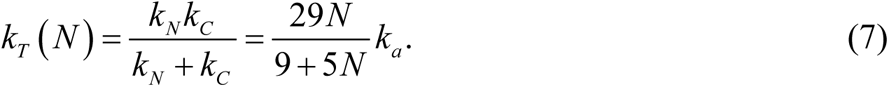

The stiffness of an isolated ARD *k_a_* can be determined by the measured stiffness *k_T_* (1) =0.7 pN/nm. The stiffnesses with multiple subunits being pulled are calculated using Eq. (7) as shown in Fig. 5a. The addition of the C-terminal contact site (CS2) does not change our estimations.

## Acknowledgments

We thank Joe Howard and Fred Sigworth for reading our manuscript and constructive discussion. This work was supported by NIH grants R35 GM131714 to Y. Z.

## Author contributions

Y. W., P. J., A. K., L. J., Y. C., Y. J., and Y. Z. designed the experiments, Y. W., P. J., A. K. performed the experiments, Y. W., P. J., A. K., Y. C., Y. J., and Y. Z. analyzed the data, and Y. W., P. J., A. K., Y. J., and Y. Z. wrote the paper.

**Competing interest**

The authors declare no interest of conflict.

