## Supplementary materials for "Nonlinear compliance of NompC gating spring and its implication in mechanotransduction"

#### Supplementary Information

**Extended Data Fig. 1. FECs of different NompC complexes revealed common unfolding intermediate states with different lifetimes.** FECs (black) of three different NompC complexes and their associated long-lived intermediate states (I1, I2, and I3). The dashed curves are the best fits of the worm-like chain model to different overlapped FEC regions. The first transition is often reversible (region marked by dashed blue rectangles).

**Extended Data Fig. 2. FECs of NompC complexes showing refolding ARDs.** Single NompC complexes were first pulled to unfold their ARDs to different intermediates (black curves) and then relaxed from these states to detect their refolding at a low force (cyan curves). The fully folded ARDs were confirmed by a second pulling round, whose FECs (green) generally overlapped the FECs from the first pulling round.

**Extended Data Fig. 3**. **Purification of AnkB ARD.** Gel filtration elution profile of the purified AnkB ARD (peak indicated by the black arrow).

**Extended Data Fig. 4**. **FECs showing parallel folding pathways of AnkB ARD**. FECs of three AnkB ARD molecules show highly parallel unfolding pathways.

**Movie 1. Cartoons showing steps of toehold-mediated strand displacement strategy to insert a single protein into pre-stretched DNA handles for single-molecule manipulation experiments.**

**Extended Data Fig. 1**

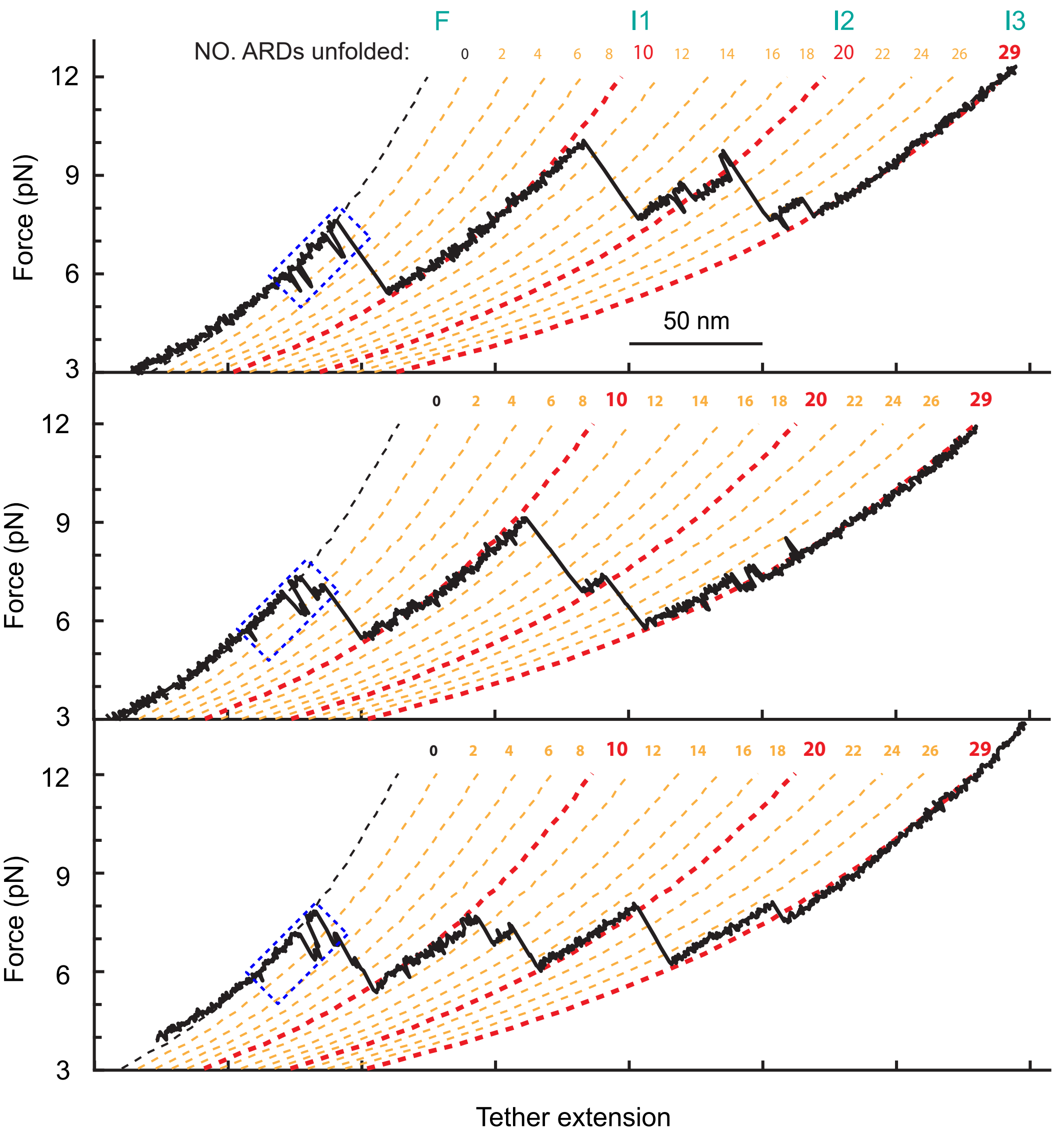

#### Extended Data Fig. 2

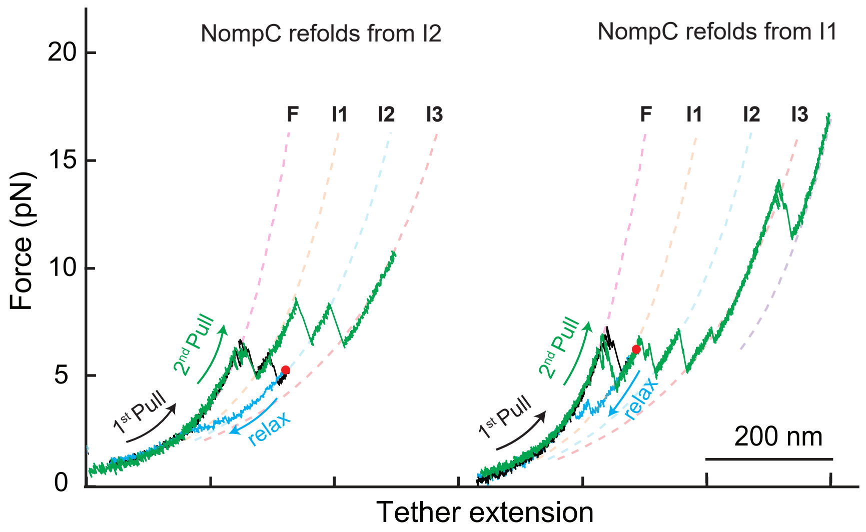

### Extended Data Fig. 3

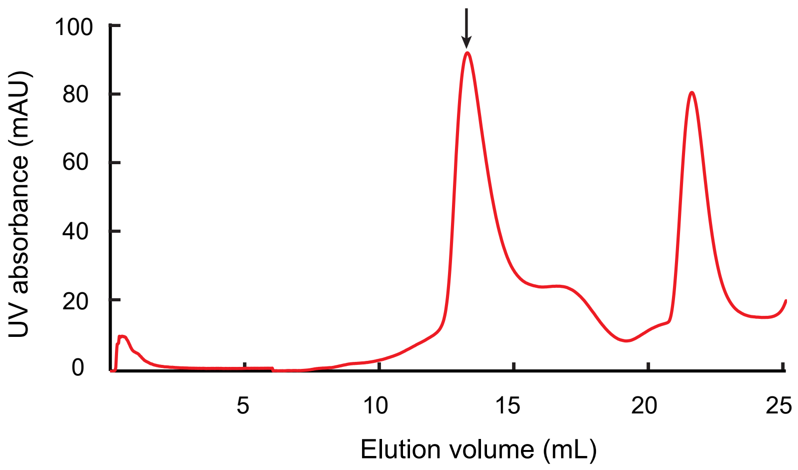

### Extended Data Fig. 4

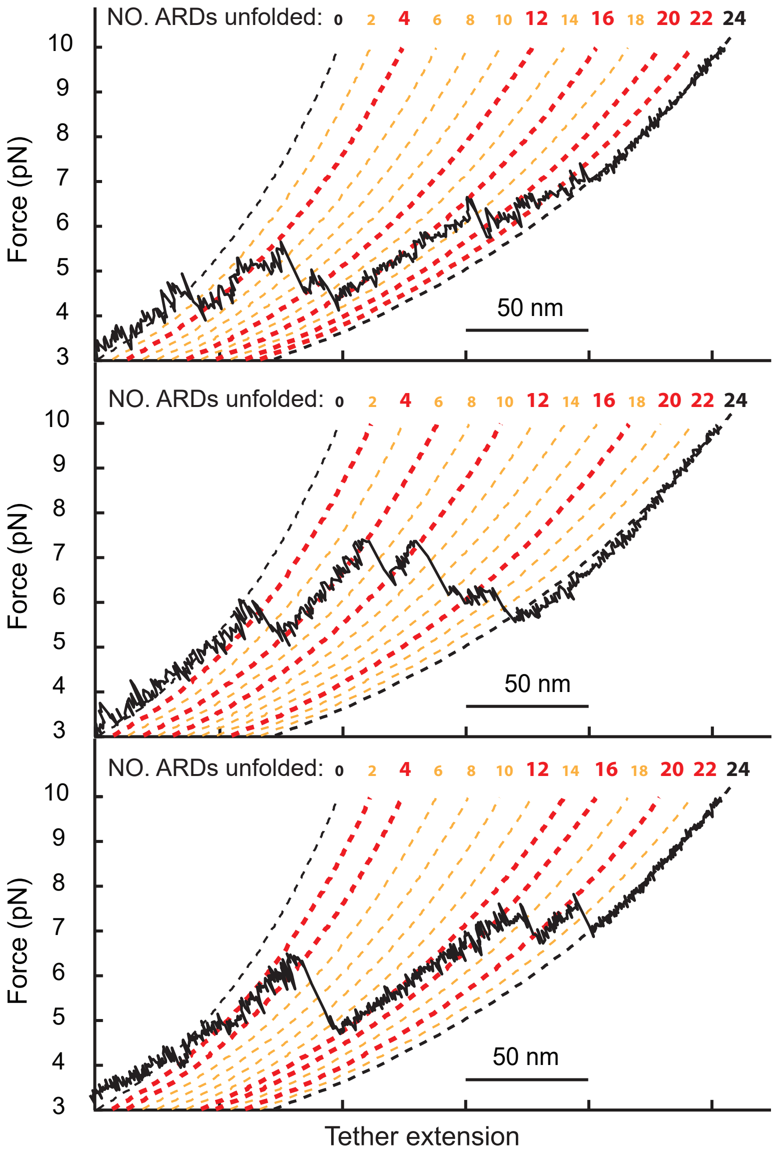
